# Quantitative physiology of non-energy-limited retentostat cultures of *Saccharomyces cerevisiae* at near-zero specific growth rates

**DOI:** 10.1101/653816

**Authors:** Yaya Liu, Anissa el Masoudi, Jack T. Pronk, Walter M. van Gulik

**Author notes:** Corresponding author: W.M. van Gulik. Royal Haskoning DHV, George Hintzenweg 85, 3009 AM Rotterdam, The Netherlands.

## Abstract

So far, the physiology of *Saccharomyces cerevisiae* at near-zero growth rates has been studied in retentostat cultures with a growth-limiting supply of the carbon and energy source. Despite its relevance in nature and industry, the near-zero growth physiology of *S. cerevisiae* under conditions where growth is limited by the supply of non-energy substrates remains largely unexplored. This study analyses the physiology of *S. cerevisiae* in aerobic chemostat and retentostat cultures grown under either ammonium or phosphate limitation. To compensate for loss of extracellular nitrogen- or phosphorus-containing compounds, establishing near-zero growth rates (μ < 0.002 h^-1^) in these retentostats required addition of low concentrations of ammonium or phosphate to reservoir media. In chemostats as well as in retentostats, strongly reduced cellular contents of the growth-limiting element (nitrogen or phosphorus) and high accumulation levels of storage carbohydrates were observed. Even at near-zero growth rates, culture viability in non-energy-limited retentostats remained above 80 % and ATP synthesis was still sufficient to maintain an adequate energy status and keep cells in a metabolic active state. Compared to similar glucose-limited retentostat cultures, the nitrogen- and phosphate-limited cultures showed a partial uncoupling of catabolism and anabolism and aerobic fermentation. The possibility to achieve stable, near-zero growth cultures of *S. cerevisiae* under nitrogen- or phosphorus-limitation offers interesting prospects for high-yield production of bio-based chemicals.

**Importance:** The yeast *Saccharomyces cerevisiae* is a commonly used microbial host for production of various bio-chemical compounds. From a physiological perspective, biosynthesis of these compounds competes with biomass formation in terms of carbon and/or energy equivalents. Fermentation processes functioning at extremely low or near-zero growth rates would prevent loss of feedstock to biomass production. Establishing *S. cerevisiae* cultures in which growth is restricted by the limited supply of a non-energy substrate could therefore have a wide range of industrial applications, but remains largely unexplored. In this work we accomplished near-zero growth of *S. cerevisiae* through limited supply of a non-energy nutrient, namely the nitrogen or phosphorus source and carried out a quantitative physiology study of the cells under these conditions. The possibility to achieve near-zero-growth *S. cerevisiae* cultures through limited supply of a non-energy nutrient may offer interesting prospects to develop novel fermentation processes for high-yield production of bio-based chemicals.

## Introduction

The yeast *Saccharomyces cerevisiae* is an established microbial host for production of a wide range of bio-chemical compounds (1, 2). Current aerobic processes for production of ATP-requiring (‘anabolic’) products are typically biphasic, with separate growth and production phases. Complete uncoupling of growth and product formation could enable a further reduction of the loss of feedstock to biomass production. In theory, such a complete uncoupling can be achieved in continuous processes performed at very low or near-zero specific growth rates. In practice, however, its implementation requires processes and microorganisms that, over prolonged periods of time, ensure a high viability and a high biomass-specific product formation rate (q_p_) in the absence of growth.

For laboratory studies near-zero specific growth rates are usually achieved in retentostats (3). A retentostat is a modification of the chemostat, in which effluent removal occurs through an internal or external filter module that causes complete biomass retention. Retentostats enable studies on microbial physiology at near-zero growth rates that are technically difficult to achieve in conventional chemostats, while their use avoids complete starvation by maintaining a constant supply of essential nutrients.

When growth in retentostat cultures is limited by the energy substrate, biomass accumulates in the reactor until the biomass-specific substrate consumption rate (q_s_) equals the energy-substrate requirement for cellular maintenance (m_s_). Aerobic and anaerobic, glucose-limited retentostat cultures of *S. cerevisiae* were shown to retain a high viability, as well as an extremely high heat-shock tolerance, over periods of several weeks (4–7). Consistent with a growth-rate-independent requirement of ATP for cellular maintenance (8), observed values of q_s_ at near-zero growth rates (μ < 0.002 h^-1^) were in good agreement with estimates of m_s_ derived from measurements in glucose-limited chemostat cultures grown at a range of specific growth rates (4, 6).

From an applied perspective, it seems illogical to apply severely energy-limited cultivation regimes for production of compounds whose synthesis from sugar requires a net input of ATP. In nature, *S. cerevisiae* seems to have primarily evolved for growth in sugar-rich environments where, instead of the energy substrate, the nitrogen source is growth limiting (9, 10). Also in industrial substrates for *S. cerevisiae* such as wine most or brewing wort, sugar is typically present in abundance, while growth becomes limited by the nitrogen source (11). As an alternative to nitrogen-limited cultivation, growth under extreme phosphate limitation may offer interesting options to uncouple growth from product formation. For example, *S. cerevisiae*, a non-oleaginous yeast, has been reported to accumulate high levels of specific fatty acids when availability of phosphate is restricted (12). Studies in exponentially growing chemostat cultures have revealed an extensive reprogramming of the yeast transcriptome, proteome and fluxome in response to nitrogen and phosphorus limitation (13–16). In addition, nitrogen- and phosphorus-limited growth of resulted in lower contents of protein and phospholipids, respectively, in yeast biomass (17, 18). In contrast to the wealth of data on the effects of different nutrient limitation regimes in actively growing cultures, information on aerobic *S. cerevisiae* cultures grown at near-zero growth rates is scarce. In anaerobic cultures, nitrogen-limited cultivation with biomass recycling has been explored to maximize ethanol yields (19, 20). Brandberg and coauthors (21), who investigated the impact of severe nitrogen limitation on ethanol production by *S. cerevisiae*, used incomplete cell recycling under anaerobic and micro-aerobic conditions.

The goal of the present study is to design and implement retentostat regimes for aerobic, nitrogen- and phosphate-limited growth of *S. cerevisiae* at near-zero specific growth rates and to use the resulting cultures for a first experimental exploration of its quantitative physiology under these scientifically interesting and industrially relevant conditions. To this end, experimental setups were tested that allowed for a smooth transition from low growth rate chemostat cultures to near-zero growth rate retentostat cultures. Metabolic fluxes, biomass composition and cellular robustness were analysed and compared with previously obtained data from glucose-limited chemostat and retentostat cultures.

## Results

### Design of carbon-excess retentostat regimes

To study the physiology of *S. cerevisiae* at near-zero growth rates under non-energy-limited conditions, retentostat regimes were designed in which growth was prevented by a severely limited supply of ammonium or phosphate. To avoid starvation, any loss of nitrogen or phosphate from such cultures, either by cell lysis or by excretion of N- or P-containing compounds from viable cells, should be compensated for. As a first approximation of the rates of N and P release by *S. cerevisiae* at near-zero growth rates, concentrations of N- and P-containing compounds were quantified in the outflow of an aerobic, glucose-limited retentostat culture. From these measurements, biomass-specific release rates of 8.1 μmol N/[g biomass]/h and 5.2 μmol P/[g biomass]/h were calculated (Supplementary Table S1). These rates were used to estimate required supply rates of ammonium and phosphate in non-growing retentostat cultures limited by either of these two nutrients. For a target biomass concentration in the retentostats of 5 g/L at a dilution rate of 0.025 h^-1^, 0.1 g/L (NH_4_)_2_SO_4_ was included in the medium feed of the ammonium-limited cultures, while 0.014 g/L KH_2_PO_4_ was used for phosphate-limited retentostat cultivation.

Aerobic growth of *S. cerevisiae* at non-limiting concentrations of glucose leads to aerobic alcoholic fermentation (22). Based on trial experiments, glucose concentrations in the influent of ammonium- and phosphate-limited retentostats were set at 120 g/L and 60 g/L, respectively. These concentrations of the growth-limiting nutrients resulted in residual glucose concentrations of ca. 15 g/L. Ethanol concentrations did not exceed 20 g/L, which is well below the value of 5 % (v/v) that has been reported to cause stress responses (23).

### Growth and viability in ammonium- and phosphate-limited retentostat cultures

Retentostat cultures were started by redirecting the effluent of steady-state ammonium- or phosphate-limited chemostat cultures, grown at a dilution rate of 0.025 h^-1^, through a membrane filter unit placed inside the reactor (see Materials and Methods). Replicate ammonium-limited retentostats were operated for 220 h with full biomass retention, after which fouling caused the membrane filters to clog. Membrane fouling was not observed in the phosphate-limited retentostats, which were operated with full biomass retention until, after 400 h, the biomass concentration had reached a stable value.

Irrespective of the nutrient limitation regime, the onset of retentostat cultivation led to a gradual increase of the biomass concentration (Fig. 1A and 1B). In ammonium-limited retentostats, the biomass concentration stabilized at ca. 14 g/L after 150 h, while stabilization in the phosphate-limited cultures at ca. 18 g/L occurred after 300 h. The increase in biomass concentration in the ammonium-limited retentostats mainly reflected an increase of the dry mass per cell, which was initially smaller than in the phosphate-limited retentostats. Conversely, the biomass increase in phosphate-limited retentostats predominantly reflected an increase of the cell number (Fig. 1C and 1D).

**Fig. 1.**
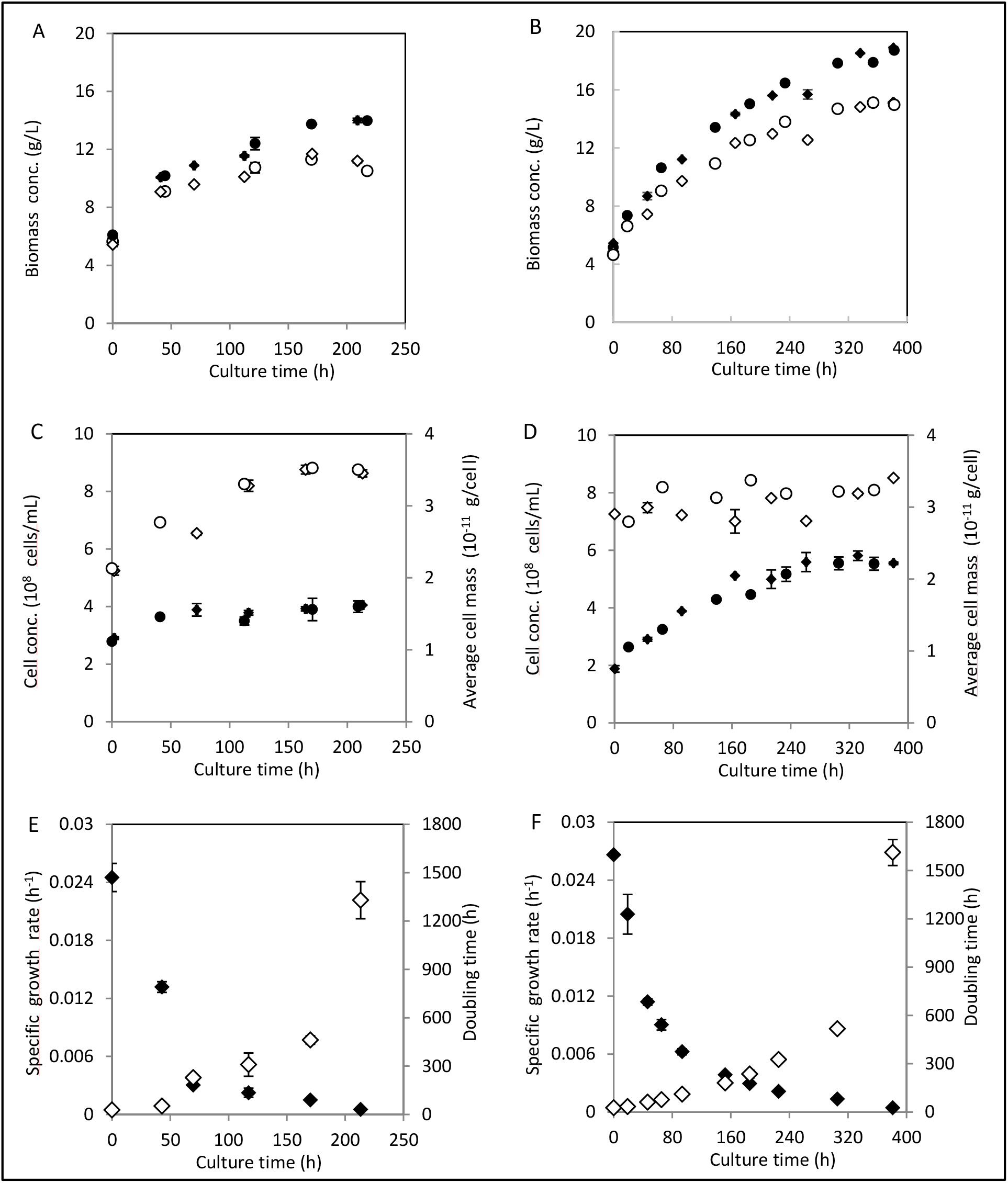
Biomass accumulation, cell counts and specific growth rates in aerobic ammonium- and phosphate-limited retentostat cultures of *S. cerevisiae* CEN.PK113-7D. Data of Fig. 1A, 1B, 1C, and 1D obtained from independent duplicate cultures are shown as circles and diamonds, and error bars indicate standard errors of analytical replicates on samples from the same culture. Data of Fig. 1C and 1D represent the averages and standard errors of measurements on duplicate retentostat cultures. A, B: Total biomass (closed symbols), viable biomass (open symbols) and percentage of viable biomass in ammonium-limited (A) and phosphate-limited (B) retentostat cultures. C, D: Cell numbers (closed symbols) and average mass per cell (open symbols) in ammonium-limited (C) and phosphate-limited (D) retentostat cultures. E, F: Specific growth rate (closed symbols) and doubling time (open symbols) in ammonium-limited (E) and phosphate-limited (F) retentostat cultures.

Culture viability was estimated by plate counts of colony-forming units (CFU) and by flow cytometry after CFDA/propidium iodide (PI) staining (Supplementary Table S2). We observed a consistently lower viability in the CFU assays than in the CFDA/PI stains. A similar difference has previously been attributed to loss of viability of retentostat-grown cells during plating (4, 6). Based on PI staining, the viability of the ammonium- and phosphate-limited retentostat cultures towards the end of the experiments did not decrease below 80 % and 90 %, respectively (Fig. 1A and 1B, Supplementary Table S2).

During retentostat cultivation, specific growth rates progressively decreased, reaching final values of 0.00056 ± 0.00010 h^-1^ and 0.00043 ± 0.00012 h^-1^ for the ammonium- and phosphate-limited cultures, respectively, corresponding to doubling times of 55 and 67 days (Fig. 1E and 1F). Based on these observations, death rates of 0.0018 ± 0.0001 h^-1^ and 0.0012 ± 0.0001 h^-1^ were calculated for prolonged ammonium- and phosphate-limited retentostat cultures, respectively. The resulting gradual decrease of culture viability partially explained the difference between the observed biomass accumulation and the targeted values in the experimental design.

### Quantitative physiology under extreme ammonium and phosphate limitation

During retentostat cultivation, the biomass-specific consumption rates of glucose and oxygen and production rates of ethanol and CO_2_ asymptotically decreased over time and stabilized after approximately 100 h in the ammonium-limited cultures and after approximately 200 h in the phosphate-limited cultures (Supplementary Fig. S1). At this stage, the specific growth rate of the cultures was lower than 0.002 h^-1^, growth stoichiometries became constant (Fig. 1E and 1F) and cells were assumed to be in a metabolic pseudo steady state. Physiological parameters obtained from the preceding, slowly growing steady-state chemostat cultures (μ = 0.025 h^-1^) and from the pseudo-steady-state, near-zero growth retentostat cultures (μ < 0.002 h^-1^) are summarized in Table 1. As anticipated, the concentrations of the limiting nutrients (ammonium or phosphate) were below the detection limit, whereas glucose concentrations were between 10 and 20 g/L in all cultures (Table 1). Carbon- and degree-of-reduction balances yielded recoveries close to 100 % (Table 1), indicating that no major metabolites had been overlooked in the analyses.

**Table 1.**
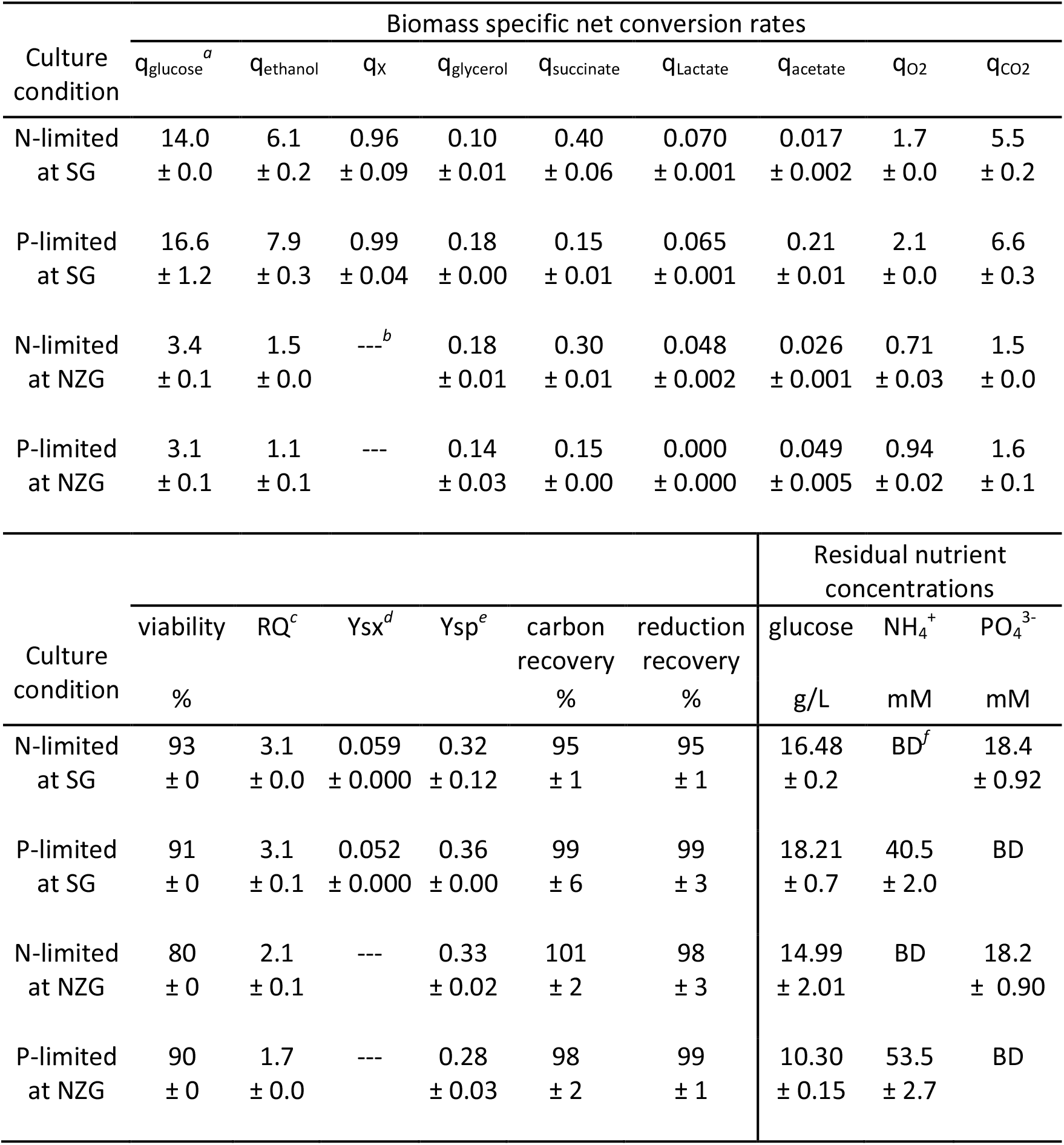
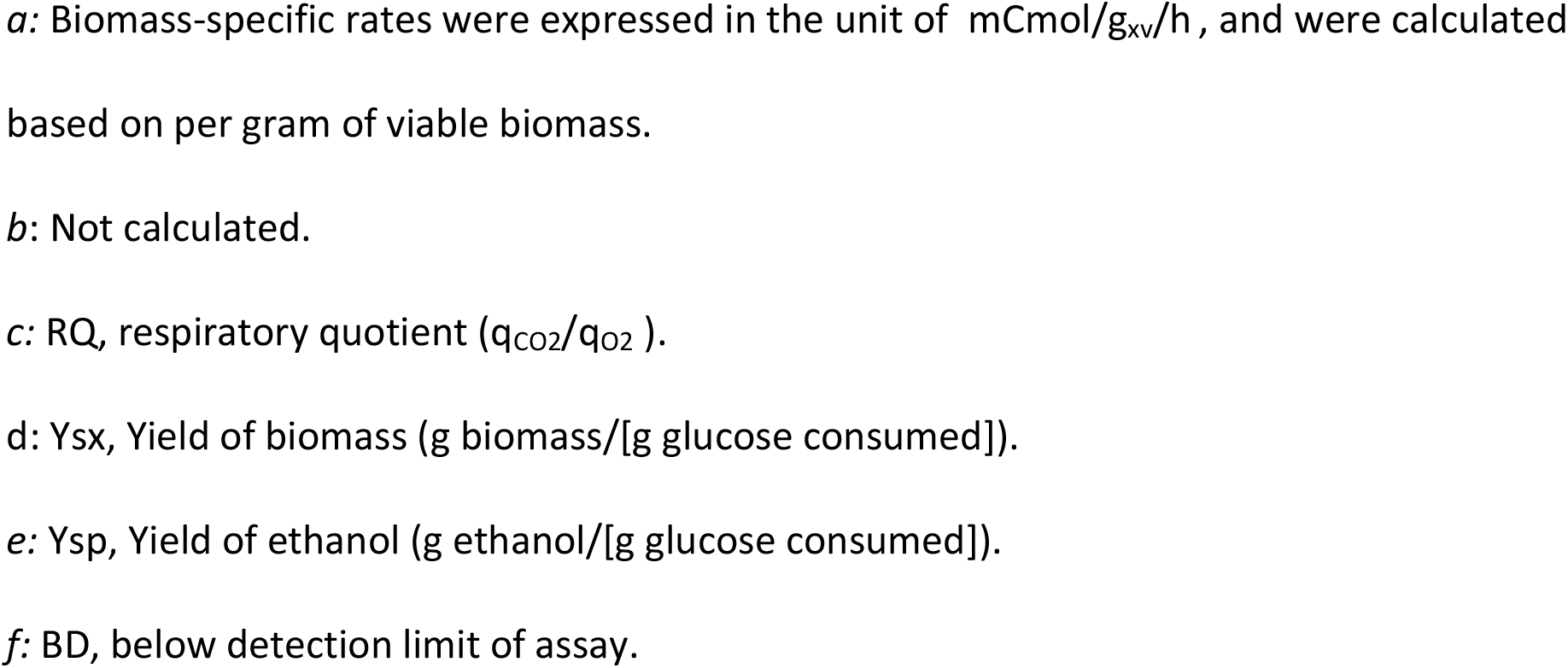
Physiological parameters of *S. cerevisiae* CEN.PK113-7D cultured in aerobic ammonium- and phosphate-limited (N- and P-limited) slow growth (SG) (μ = 0.025 h^-1^) steady-state chemostats and near-zero growth (NZG) (μ < 0.002 h^-1^) pseudo-steady-state retentostats. Data represent averages, with their standard errors, calculated from multiple measurements obtained from duplicate experiments.

In the slow-growing (μ = 0.025 h^-1^) chemostat cultures the biomass-specific rates of glucose and oxygen consumption as well as ethanol and carbon dioxide production, were consistently higher in the phosphate-limited cultures than in the ammonium-limited cultures (Table 1). In line with these observations, the phosphate-limited cultures showed a lower biomass yield and higher ethanol yield on glucose. Respiratory quotients (RQ, ratio of CO_2_ production and O_2_ consumption rate) were identical for the two nutrient limitation regimes, indicating that the difference in biomass yield of the chemostat cultures was not caused by different contributions of respiratory and fermentative metabolism. Furthermore, the sum of the specific production rates of the four minor byproducts (glycerol, succinate, lactate and acetate), which accounted for less than 4 % of the consumed glucose, were not significantly different for the two limitation regimes and were also not responsible for the observed difference in biomass yield.

In the pseudo-steady-state near-zero growth retentostat cultures, the observed ethanol yields on glucose (Table1) were respectively 71 % and 53 % of the theoretical maximum (0.51 g ethanol/[g glucose]) for the ammonium- and phosphate-limited regimes. Consistent with this observation, significant oxygen consumption occurred in these cultures and their RQ values were significantly lower than those of the preceding chemostat cultures. For the phosphate-limited cultures the difference was most pronounced. These observations indicate that near-zero growth achieved by phosphate limitation leads to a more respiratory metabolism than was observed in the preceding slowly growing, phosphate-limited chemostats. Formation of byproducts accounted for 16 % and 11 % of the supplied glucose in the ammonium- and phosphate-limited near-zero growth cultures, respectively. Glycerol and succinate were the main contributors, with succinate accounting for 9 % of the consumed glucose in the ammonium-limited culture.

### Biomass composition under extreme ammonium and phosphate limitation

To analyse the impact of extreme ammonium and phosphate limitation on biomass composition, biomass samples from slow growing, steady-state chemostat cultures and from near-zero growth rate pseudo-steady-state retentostat cultures were analysed for their elemental and macromolecular compositions (Table 2). In the chemostat cultures as well as in the retentostat cultures, the content of the growth-limiting element in the biomass was strongly reduced relative to that of the culture grown under the other nutrient limitation (Table 2). This difference was even more pronounced in the retentostat cultures than in the preceding chemostat cultures. The nitrogen content of biomass from ammonium-limited retentostat cultures was ca. 2-fold lower than that of the corresponding phosphate-limited retentostats, while the phosphorus content of biomass from the phosphate-limited retentostats was 3.5-fold lower than that of biomass from the ammonium-limited retentostats. Both in phosphate-limited chemostats and retentostats, a low phosphorus content was accompanied by a 2-3 fold higher sulfur content than in the corresponding ammonium-limited cultures. The increased sulfur content in phosphate-limited cultures may be due to sulfate uptake by high-affinity phosphate transporters (14). Compared with glucose-limited chemostat cultures of the same *S. cerevisiae* strain at a similar dilution rate (D= 0.022 h^-1^, Table 2), the biomass protein content and the total nitrogen content of cells grown in the ammonium-limited chemostat cultures were over 60 % and 50 % lower, respectively. Similarly, in the phosphate-limited chemostat cultures, the phosphorus content of the biomass was ca. 50 % lower.

**Table 2.** Biomass elemental compositions of *S. cerevisiae* CEN.PK113-7D cultured in aerobic ammonium- and phosphate-limited (N- and P-limited) slow growth (SG) (μ = 0.025 h^-1^) steady-state chemostats and near-zero growth (NZG) (μ < 0.002 h^-1^) pseudo-steady-state retentostats. Data represent averages, with standard errors, of measurements from duplicate cultures and are compared with published values from aerobic glucose-limited (C-limited) chemostat culture of the same strain (49).

| Culture condition | $\mu$<br>$\text{h}^{-1}$ | $f_C$<br>% | $f_H$<br>% | $f_N$<br>% | $f_O$<br>% | $f_P$<br>% | $f_S$<br>% | sum<br>% | C-mol weight<br>g/L |
| --- | --- | --- | --- | --- | --- | --- | --- | --- | --- |
| N-limited | 0.025 | 47.0<br>$\pm 0.0$ | 7.2<br>$\pm 0.0$ | 3.5<br>$\pm 0.1$ | 39.5<br>$\pm 0.4$ | 1.3<br>$\pm 0.0$ | 0.16<br>$\pm 0.01$ | 98<br>$\pm 0$ | 25.56<br>$\pm 0.00$ |
| P-limited | 0.025 | 44.0<br>$\pm 0.2$ | 7.0<br>$\pm 0.1$ | 5.3<br>$\pm 0.0$ | 36.9<br>$\pm 0.3$ | 0.50<br>$\pm 0.01$ | 0.39<br>$\pm 0.03$ | 94<br>$\pm 0$ | 24.27<br>$\pm 0.25$ |
| N-limited | < 0.002 | 49.5<br>$\pm 0.4$ | 7.6<br>$\pm 0.0$ | 2.5<br>$\pm 0.0$ | 39.5<br>$\pm 1.1$ | 1.1<br>$\pm 0.0$ | 0.11<br>$\pm 0.01$ | 100<br>$\pm 1$ | 27.33<br>$\pm 0.22$ |
| P-limited | < 0.002 | 47.3<br>$\pm 0.1$ | 7.2<br>$\pm 0.0$ | 4.6<br>$\pm 0.1$ | 37.4<br>$\pm 0.0$ | 0.29<br>$\pm 0.00$ | 0.27<br>$\pm 0.01$ | 97<br>$\pm 0$ | 25.42<br>$\pm 0.08$ |
| C-limited | 0.022 | 45.6 | 6.8 | 6.6 | 37.3 | 1.0 | 0.22 | 97 | 26.4 |

Consistent with their low nitrogen content, ammonium-limited chemostat and retentostat cultures showed a ca. 2.5-fold lower biomass protein content than the corresponding phosphate-limited cultures, with the lowest protein content (9.6 %) measured in the ammonium-limited retentostats (Fig. 2A). Conversely, glycogen contents were higher (5.8 fold in chemostats and 1.8 fold in retentostats) in ammonium-limited cultures than in phosphate-limited cultures, while trehalose contents were only 30-40 % higher in the ammonium-limited cultures (Fig. 2B). When analysed throughout the retentostat experiments, glycogen contents in the ammonium-limited cultures remained consistently high, while they increased with declining specific growth rate in the phosphate-limited cultures (Fig. 2C). For both nutrient limitation regimes, the trehalose content reached a maximum at a specific growth rate of ca. 0.01 h^-1^ (Fig. 2D).

**Fig. 2.**
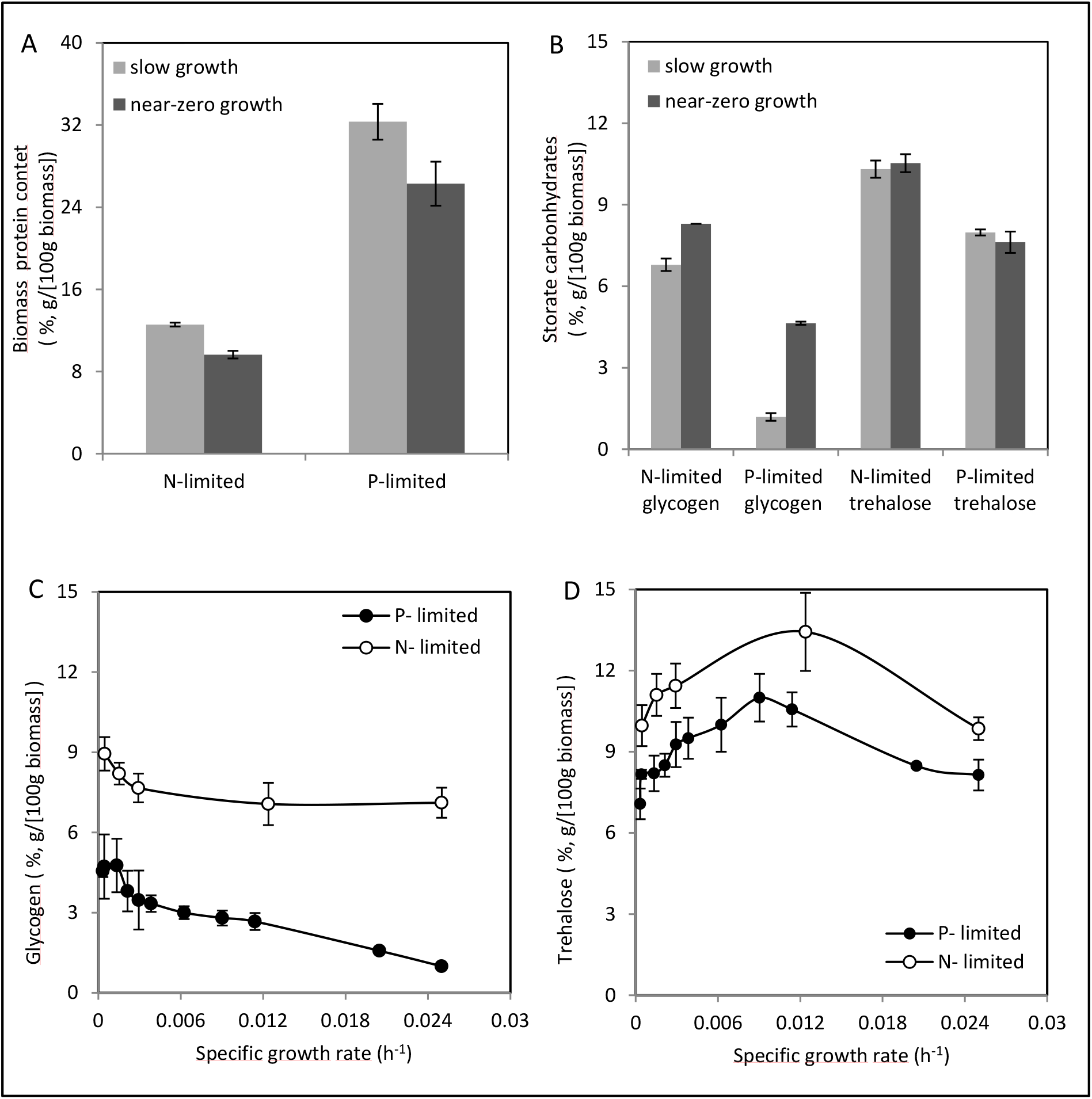
Biomass protein and storage carbohydrates (glycogen and trehalose) contents in aerobic ammonium- and phosphate-limited (N- and P-limited) cultures of *S. cerevisiae* CEN.PK113-7D. Data represent the averages and standard errors of multiple measurements on duplicate cultures. A, B: Biomass protein (A) and storage carbohydrates(glycogen and trehalose) (B). Samples were withdrawn from the steady-state, slow growth (μ = 0.025 h^-1^) chemostat cultures, and the pseudo-steady-state, near-zero growth (μ < 0.002 h^-1^) retentostat cultures. C, D: Glycogen (C) and trehalose (D) contents vs. the specific growth rate in the prolonged retentostat cultures.

### Metabolic flux analysis

To further investigate the physiological differences between extreme ammonium and phosphate limitation, metabolic flux analysis was performed for both the slow growing, steady-state chemostat cultures (μ = 0.025 h^-1^) and near-zero growth, pseudo-steady-state retentostat cultures (μ < 0.002 h^-1^) (Fig. 3, Supplementary Table S3). At a specific growth rate of 0.025 h^-1^, fluxes through the glycolysis, tricarboxylic acid cycle (TCA cycle) and pyruvate branch point were consistently higher in the phosphate-limited cultures than in the ammonium-limited cultures. This observation indicated a higher contribution of catabolism in the phosphate-limited cultures. Assuming a P/O ratio of 1 (24), biomass-specific rates of ATP turnover were ca. 1.3-folder higher in the phosphate-limitated chemostat cultures than in the corresponding ammonium-limited cultures (Fig. 3).

**Fig. 3.**
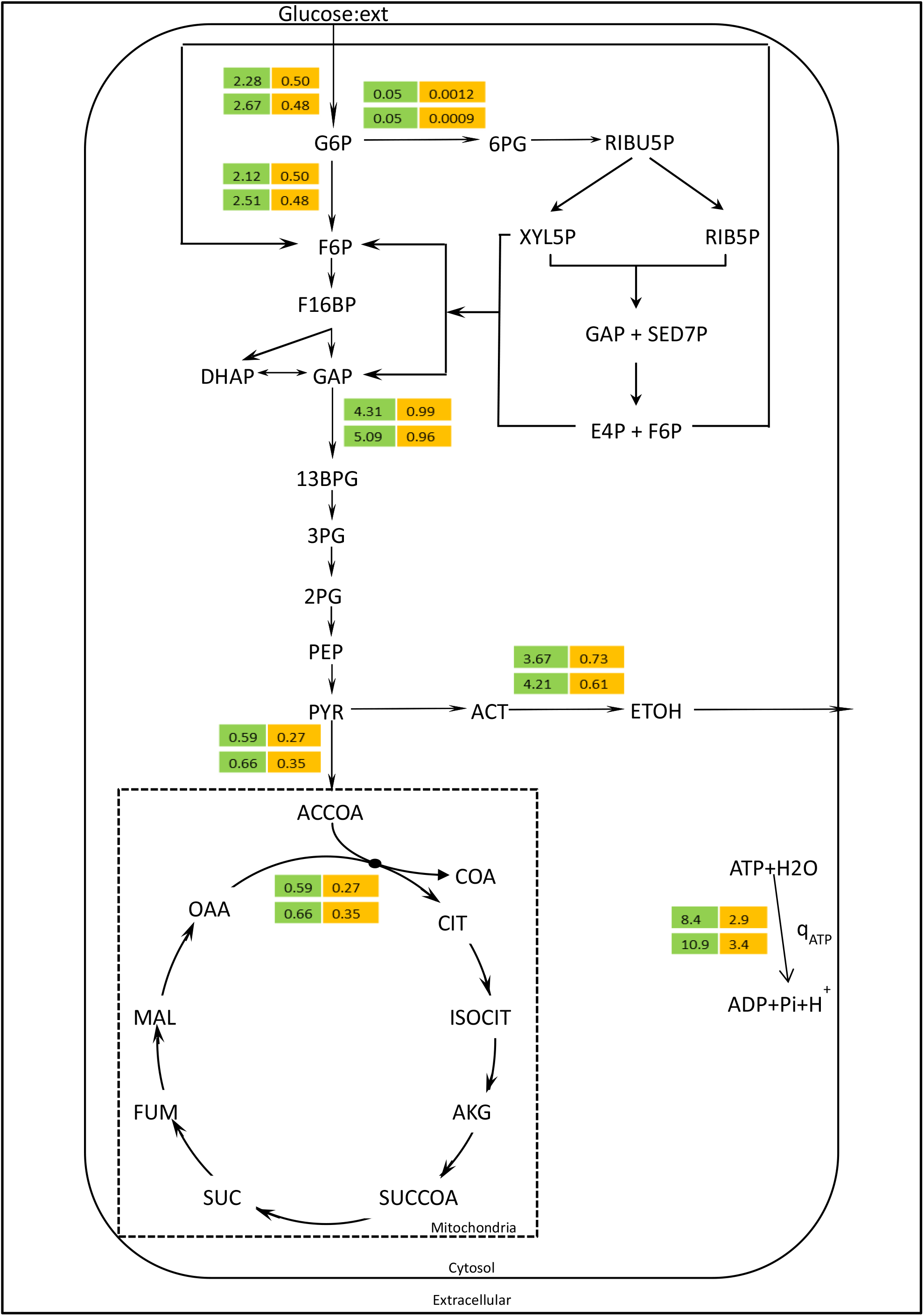
Metabolic flux analysis in aerobic ammonium- and phosphate-limited (N- and P-limited) cultures of *S. cerevisiae* CEN.PK113-7D. Flux values present the steady-state, slow growth (SG) (μ = 0.025 h^-1^) chemostat cultures (numbers on green background), and the pseudo-steady-state, near-zero growth (NZG) (μ < 0.002 h^-1^) retentostat cultures (numbers on orange background). Data are expressed in millmoles of per gram viable biomass per hour and represent the averages of duplicate cultures. Complete flux analysis values and standard errors were presented in Supplementary Table S3.

In the retentostats, fluxes through the pentose-phosphate pathway (PPP) were extremely low, which is consistent with the strictly assimilatory role of this central metabolic pathway in *S. cerevisiae* (25). The glycolytic flux was nearly identical for the two nutrient limitations. Conversely, distribution of pyruvate over alcoholic fermentation and TCA cycle were different. Consistent with their lower RQ, phosphate-limited retentostat cultures channeled a higher fraction of the pyruvate into the TCA cycle than the ammonium limited retentostat cultures. Estimated non-growth-associated ATP consumption was higher in the phosphate-limited retentostats (3.4 ± 0.2 mmol ATP/[g viable biomass]/h) than in the ammonium-limited retentostats (2.9 ± 0.1 mmol ATP/[g viable biomass]/h) (Fig. 3).

### Energetics under extreme ammonium and phosphate limitation

Nitrogen and phosphate limitation can both be characterized as non-energy-limited cultivation regimes. However, because phosphate plays a vital role in cellular energy metabolism and energy status, the intracellular nucleotide levels (ATP, ADP and AMP) and corresponding adenylate energy charge and ATP/ADP ratios were quantified for both chemostat and retentostat conditions (Fig. 4). Intracellular levels of all three adenine nucleotides were consistently higher in the chemostats than in the retentostats. Comparing these two limitations, both in slow-growth and near-zero growth cultures, intracellular ATP and AMP levels were consistently lower under phosphate limitation than under ammonium limitation. In addition, phosphate-limited near-zero growth cultures also showed ca. 40 % lower ADP levels than the corresponding ammonium-limited cultures, while ADP levels were identical in phosphate- and ammonium-limited, slow-growing chemostat cultures (Fig. 4A). Neither the ATP/ADP ratios nor the energy charge in the retentostat cultures differed from those in the corresponding slow-growing chemostat cultures (Fig. 4B and 4C). However, ATP/ADP ratios in the phosphate-limited cultures were 30–35 % lower than in the corresponding ammonium-limited cultures. A similar, less pronounced difference was observed for the adenylate energy charge. These results show that phosphate limitation indeed significantly affected cellular energy status.

**Fig. 4.**
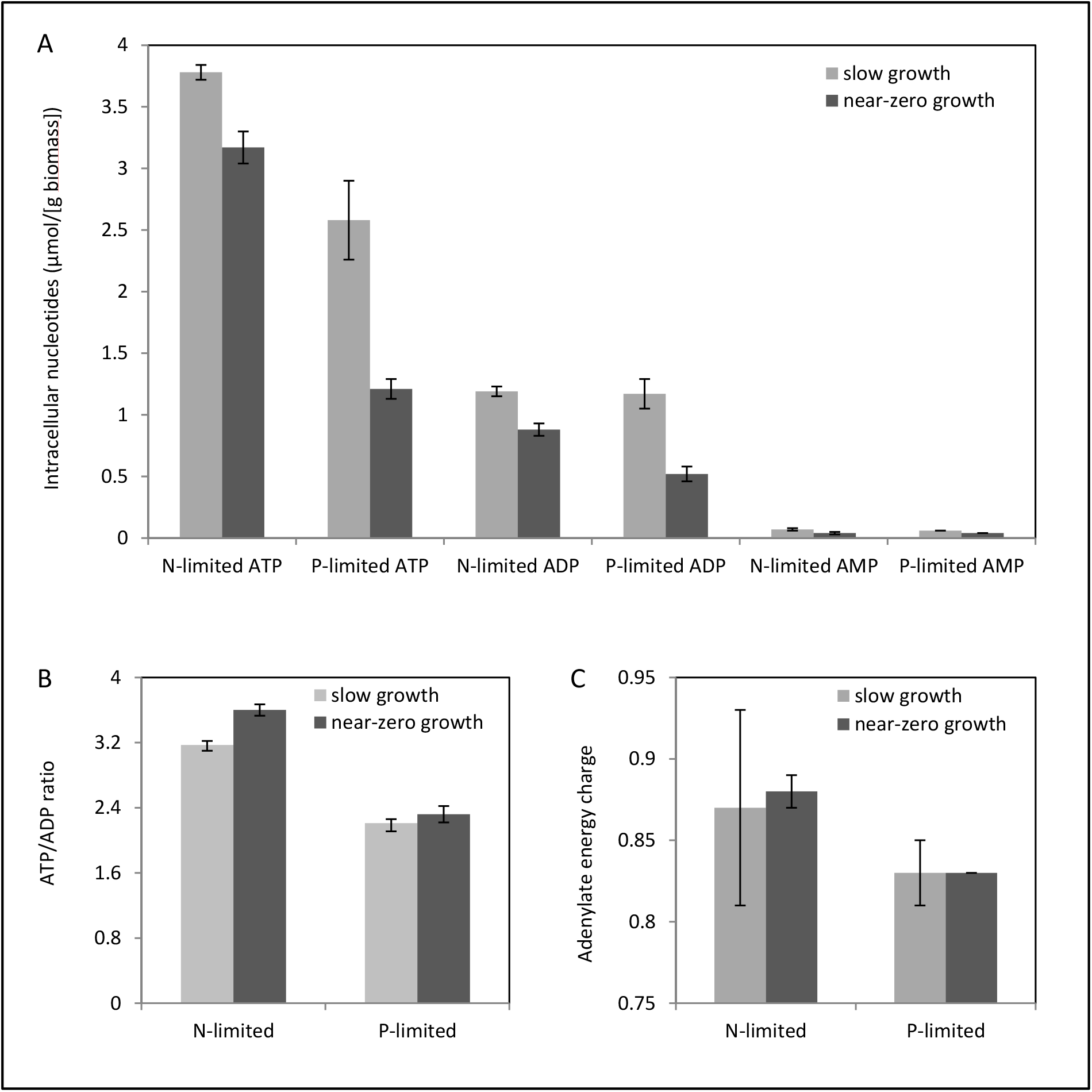
Intracellular adenosine phosphate concentrations (3A), ATP/ADP ratio (3B) and energy charge (3C) in aerobic ammonium- and phosphate-limited (N- and P-limited) cultures of *S. cerevisiae* CEN.PK113-7D. Data represent the averages and standard errors of multiple measurements from duplicate cultures. Samples were withdrawn from the steady-state, slow growth (μ = 0.025 h^-1^) chemostat cultures, and the pseudo-steady-state, near-zero growth (μ < 0.002 h^-1^) retentostat cultures.

## Discussion

### Prolonged near-zero growth of *S. cerevisiae* under non-energy-limited conditions

Retentostat cultivation of heterotrophic microorganisms typically involves a constant, growth-limiting supply rate of the carbon and energy substrate (3). The amount of viable biomass in such energy-limited retentostats asymptotically increases to a constant value, while the specific growth rate asymptotically approaches zero. In the resulting pseudo steady states, biomass-specific substrate supply rates closely match cellular maintenance-energy requirements (3). The retentostat regimes explored in this study, in which growth was restricted by supply of the nitrogen or phosphorus source, represented a fundamentally different scenario. While biomass also asymptotically increased to a constant value, the corresponding constant biomass-specific ammonium or phosphate consumption rates were not related to maintenance-energy metabolism. Instead, they represented release of nitrogen- or phosphorus-containing compounds, which were removed via the cell-free effluent.

Excretion of nitrogen- or phosphorus-containing compounds by severely ammonium- or phosphate-limited yeast cultures appears counter intuitive. Instead, release of these compounds probably occurs by cell death and/or lysis. *S. cerevisiae* can express a range of specific and non-specific amino acid permeases (26), while di- and tri-peptides can be imported by Prt2p (27). Presence of amino acids in culture supernatants is therefore likely to reflect the kinetics of such transporters, rather than a complete inability for amino-acid reconsumption by viable cells. Consistent with this hypothesis, extracellular concentrations of amino acids in the ammonium-limited retentostats were lower than the K_m_ values of the corresponding high-affinity *S. cerevisiae* amino-acid permeases (Supplementary Table S4). Biomass concentrations in the ammonium- and phosphate-limited retentostats reached values that were approximately 3-fold higher than the target value of 5 g/L on which design of growth media and operating conditions were based. This difference could only partially be attributed to accumulation of non-viable biomass. In addition, strongly reduced contents of the growth limiting element in the retentostat-grown biomass could explain this large discrepancy to a large extent.

As previously reported for glucose-limited cultures (21), ammonium- and phosphate-limited cultivation of *S. cerevisiae* at low to near-zero growth rates led to increased intracellular levels of glycogen and trehalose. This observation confirms that glycogen and trehalose accumulation is a universal physiological response of *S. cerevisiae* at near-zero growth conditions. Also in faster growing chemostat cultures, nitrogen limitation has been shown to lead to higher storage carbohydrate levels than other nutrient-limitation regimes (28). Intracellular reserves of glycogen and trehalose enable survival during carbon and energy source starvation and can fuel cell cycle progression under carbon- and energy-source limitation (29). Additionally, upregulation of genes involved in synthesis, metabolism and degradation of trehalose has been implicated in the extreme heat-shock tolerance of glucose-limited retentostat cultures of *S. cerevisiae* (6, 30).

### Energy metabolism of *S. cerevisiae* under extreme ammonium and phosphate limitation

Despite strongly reduced phosphate content and low intracellular levels of adenosine nucleotides, the adenylate energy charge of 0.83 of the phosphate-limited chemostat and retentostat cultures was within the normal physiological range of 0.7 to 0.95 (31). Also the adenylate energy charge of 0.88 for the corresponding ammonium-limited cultures indicated that cells were able to maintain their energy status under extreme nutrient restriction. Consistent with the well-known tendency of *S. cerevisiae* to exhibit aerobic alcoholic fermentation when exposed to excess glucose (22), respiratory quotients (RQs) of all ammonium- and phosphate-limited cultures were above 1. RQ values were lowest at near-zero growth rates (Supplementary Table S5), indicating that the contribution of fermentative metabolism decreased with decreasing specific growth rate. Even though *S. cerevisiae* has a low P/O ratio, respiratory catabolism of glucose yields much more ATP than fermentation (32). However, it maximum rate of fermentative ATP generation is approximately 2-fold higher than its maximum rate of respiratory ATP generation (33). These observations underlie a rate/yield trade-off hypothesis, according to which ATP can either be produced fast (but with a low efficiency) or efficiently (but at a lower maximum rate) (34). The shift towards a more respiratory metabolism in the near-zero growth rate retentostat cultures is entirely in line with this hypothesis.

Non-growth associated rates of ATP turnover in the aerobic, non-energy-limited cultures were significantly higher than maintenance-energy requirements estimated from aerobic and anaerobic energy-limited retentostat studies with the same *S. cerevisiae* strain (supplementary Fig. S3). While a similar uncoupling of anabolic energy demand and catabolic energy conservation has been reported for nitrogen-limited chemostat cultures, the underlying mechanism has not been elucidated (16, 21, 35–38). Quantification of the *in vivo* cytosolic concentrations of ammonium and ammonia recently showed that, in ammonium-limited chemostat cultures of *S. cerevisiae* grown at pH 5, cytosolic ammonia concentrations exceeded extracellular concentrations (39). Diffusion of ammonia from the cells, combined with reuptake of ammonium cation by the high-affinity uniporter Mep2 (40) and expulsion of its associated proton by the plasma-membrane H+-ATPase Pma1 could lead to a futile cycle.

Extreme phosphate-limited growth of *S. cerevisiae* induces expression of *PHO84*, which encodes a high-affinity phosphate/proton symporter and vacuolar synthesis of inorganic polyphosphate(41). By acting as a phosphorus sink, polyphosphate sustains phosphate uptake at low extracellular concentrations (41, 42). Its synthesis in yeast requires activity of the vacuolar H+-ATPase (V-ATPase) to maintain a proton-motive force across the vacuolar membrane (41). Although high-affinity phosphate import and subsequent vacuolar polyphosphate synthesis must have resulted in increased ATP requirements, these are negligible compared to the observed non-growth associated ATP requirements in the phosphate-limited retentostat cultures. Unless very significant turnover of the polyphosphate pool has occurred, these additional ATP requirements are likely to have been caused by other, yet unknown processes.

### Possible application of severe ammonium or phosphate limitation for industrial processes

Metabolic engineering of *S. cerevisiae* has enabled the production of a wide range of compounds whose biosynthesis from sugars requires a net input of ATP (43). The specific rate of formation of such ‘anabolic’ products is determined by the capacities and regulation of the enzymes of the product pathway and connected primary metabolic pathways, as well as by the continuous (re)generation of cofactors such as NAD(P)H, Coenzyme A and ATP. To optimize yields of such products, allocation of sugar to growth should be minimized. At the same time, ATP availability should not limit product formation rates. Theoretically, these goals can be reconciled by near-zero-growth-rate cultivation under non-energy-limited conditions. This study shows that, under ammonium limitation as well as under phosphate limitation, glucose-sufficient, near-zero-growth retentostat cultures of a laboratory strain of *S. cerevisiae* is able to maintain a normal energy charge and showed only a modest loss of culture viability. The extremely low protein content of biomass grown in the nitrogen-limited retentostats is likely to represent a disadvantage for high-level expression of heterologous product pathways. Moreover, nitrogen limitation is intrinsically poorly suited for production of proteins and other nitrogen-containing compounds. Extreme phosphate limitation did not affect biomass protein levels. However, relative to glucose-limited retentostats, both the ammonium- and phosphate-limited cultures showed increased rates of non-growth associated ATP dissipation. This increase is undesirable in industrial contexts, as the resulting increased rate of sugar dissimilation would go at the expense of the product yield. Future research should therefore aim at identifying the causes of non-growth associated ATP dissipation and on their elimination, either by alternative nutrient limitation regimes, by strain engineering or by alternative approaches to restrict cell division.

## Materials and methods

### Yeast strain and media

The prototrophic, haploid yeast strain *Saccharomyces cerevisiae* CENPK 113-7D was used in this study (44). Working stocks were obtained by cultivation in YPD medium (10 g/L Bacto yeast extract, 20 g/L Bacto peptone and 20 g/L D-glucose). After addition of 30 % (v/v) glycerol, culture aliquots were stored in sterilized Eppendorf tubes at −80°C.

Ammonium- and phosphate-limited (N- and P-limited) pre-culture and batch culture media were prepared as described by Boer (16). For N-limited batch cultivation, the medium contained the following components: 1.0 g of (NH_4_)_2_SO_4_, 5.3 g of K_2_SO_4_, 3.0 g of KH_2_PO_4_, 0.5 g of MgSO_4__7H_2_O, and 59 g of glucose per liter. For P-limited batch cultivation, the medium contained 5.0 g of (NH_4_)2SO_4_, 1.9 g of K_2_SO_4_, 0.12 g of KH_2_PO_4_, 0.5 g of MgSO_4__7H_2_O, and 59 g of glucose per liter. In addition, 1 mL/L trace element solution, 1 mL/L vitamin solution and 0.2 g/L Pluronic 6100 PE antifoaming agent (BASF, Ludwigshafen, Germany) were added. Trace element and vitamin solutions were prepared as described by Verduyn (45). The compositions of media for N- and P-limited chemostat cultivation were as described above, except that the glucose concentration was increased to 120 g/L. For N-limited retentostat cultivation, the (NH_4_)2SO_4_ concentration in the medium feed was decreased to 0.1 g/L and the glucose concentration was 60 g/L. To maintain the same sulfur concentration, the K_2_SO_4_ concentration was increased to 6.46 g/L, the concentrations of the other compounds were the same as in the chemostat medium. For P-limited retentostat cultivation, the KH_2_PO_4_ concentration was lowered to 0.014 g/L and the glucose concentration was 60 g/L.

### Bioreactor set up

Bench-scale, turbine-stirred 7 L bioreactors (Applikon, Delft, The Netherlands) equipped with a single six-bladed Rushton turbine impeller with a diameter of 85 mm, were used in this study. The working volume was controlled at 5 L by placing the bioreactor on an electronic balance (Mettler Toledo, Columbus, Ohio, USA). During continuous cultivation, effluent was removed with a peristaltic pump to an effluent vessel, which was placed on an electronic balance for measurement of the dilution rate (D = 0.025 h^-1^). The culture temperature was maintained at 30 ± 0.1°C and the stirrer speed at 500 rpm. Aerobic conditions were maintained by sparging 0.5 vvm compressed air, controlled by a mass flow controller (Brooks 5850 TR, Hatfield, PA, USA). The dissolved oxygen concentration was measured on-line with a DO sensor (Mettler-Toledo GmbH, Greinfensee, Switzerland) and remained above 30 % of air saturation in all experiments. Culture pH was controlled at 5.00 ± 0.05 by automated addition of either 2 M KOH or 2 M H_2_SO_4_, using a Biostat Bplus controller (Sartorius BBI Systems, Melsungen, Germany). Exhaust gas was cooled to 4°C by an in-line condenser and dried by a Nafion dryer (Permapure, Toms River, USA) before entering a combined paramagnetic/infrared NGA 2000 off-gas analyzer (Rosemount Analytical, Anaheim, USA) for analysis of O_2_ and CO_2_ concentrations. Off-gas data were acquired with MFCS/win 3.0 software (Sartorius BBI Systems, Melsungen, Germany).

### Pre-culture, batch, chemostat and retentostat cultures

Pre-cultures, grown in 500 mL shake flasks containing 200 mL medium, were inoculated with 2 mL of stock culture and grown at 30°C and at 200 rpm for 8 h in a B Braun Certomat BS-1 incubator (Sartorius, Melsungen, Germany). Bioreactor batch cultures were started by transferring 400 mL of preculture to a bioreactor containing 4.6 L of medium. After approximately 24 h of batch cultivation, a sharp decrease of the CO_2_ concentration in the off-gas and a corresponding increase of the dissolved oxygen concentration indicated that ammonium or phosphate was depleted. The bioreactors were then switched to chemostat cultivation mode and operated at a dilution rate of 0.025 h^-1^. Steady-state was assumed to be achieved after 5 volume changes, in which stable (less than 3 % difference over 2 volume changes) off-gas CO_2_ and O_2_ concentrations, culture dry weight and cell counts were observed. At that stage, bioreactors were switched from chemostat to retentostat mode by redirecting the culture effluent through a filtration probe assembly (Applikon, Delft, The Netherlands). Each probe was fitted with a 0.22 μ*m* tubular micro-filtration polypropylene membrane (TRACE Analytics, Brunswick, Germany). Because of the limited flow rate capacity of each filter, four filtration probes were installed in each bioreactor. Before mounting on the filtration probe and autoclaving, membranes were hydrophilized overnight in 70 % (v/v) isopropanol.

To avoid a sudden decrease of substrate concentrations during the switch from chemostat to retentostat mode, a gradual transition from chemostat to retentostat medium was accomplished by using two feed pumps. The resulting time-dependent concentrations of glucose and of the growth-limiting nutrient ((NH_4_)_2_SO_4_ or KH_2_PO_4_) in the medium are described by the following equation:

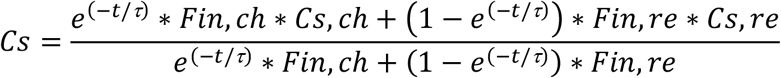

In this equation, τ is the time constant for the transition which was set to a value of 16.67 h. C_s,ch_, C_s,re_, F_in,ch_, F_in,re_, correspond to the nutrient concentrations in the chemostat and retentostat media and the feed rates from the corresponding medium reservoirs, respectively. Profiles of the resulting concentrations of the limited nutrient and of glucose in the retentostat feed media during the transition are provided in Supplementary Fig. S3. The actual medium feed rates during the chemostat and retentostat phases for each experiment were calculated from the weight increase of the effluent vessels and the addition rates of base.

### Biomass and viability assays

Culture dry weight assays were carried out through a filtration, washing and drying procedure as described previously (46). Total cell counts were quantified with a Z2 Coulter counter (50 μm aperture, Beckman, Fullerton, CA). Cell viabilities were determined through a FungaLight™ Yeast CFDA, AM/Propidium Iodide Vitality Kit (a cellular membrane integrity indicator) by flow cytometry and colony-forming-unit counts (6).

### Quantification of (by)products and residual substrates

Cell-free effluent samples were harvested from a sample port connected to the retentostat filters, immediately frozen in liquid nitrogen and stored at −80 °C until analysis. Effluent concentrations of glucose, ethanol and by-products (glycerol, lactate, acetate, and succinate) were quantified with HPLC using a Bio-Rad HPX-87H 300 column (7.8 mm). The column was eluted with phosphoric acid (1.5 mM, 0.6 mL/min). The detection was performed with a refractometer (Walters 2414) and a UV dector (Walters 484, 210 nm). Concentrations of ammonium and phosphate were quantified with an ammonium cuvette test (0.02-2.5 mg/L NH_4_^+^) and a phosphate trace cuvette test (0.03-1.5 mg/L PO4^3-^), respectively (Hach Lange GmbH, Düsseldorf, Germany).

### Balances and rate calculations

Biomass-specific glucose and oxygen consumption rates, and biomass-specific production rates of ethanol, carbon dioxide and by-products were calculated based on primary measurements of substrates/products concentration and flow rates in gas and liquid phases. Data reconciliation was performed as described previously (47). The consistencies of the thus obtained rates were evaluated by calculation of carbon and degree of reduction recoveries. Ethanol evaporation via the off-gas of the reactor was quantified as described previously (48) and was taken into account in calculation of ethanol production rates. Calculation of specific growth rates and doubling times in retentostat cultures was performed as described previously (4).

### Analysis of biomass composition

Around 250 mg of lyophilized biomass was used to determine the elemental (C, H, N, O, P, S) composition through complete combustion and subsequent gas analysis (carbon dioxide, water vapour and nitrogen mass fractions), gas chromatography (oxygen) and ICP-MS (phosphorus and sulphur) (Energy Research Centre, Petten, The Netherlands). Biomass protein was quantified with the Biuret method as described previously (49). The trehalose content of the biomass was directly quantified by GC-MS/MS (50) in intracellular metabolite samples prepared as described below. Glycogen content was quantified through an enzymatic hydrolysis method (6).

### Quantification of intracellular metabolites

A rapid sampling device connected to the bioreactor was used to rapidly withdraw broth samples for intracellular metabolite measurements (51). Approximately 1.2 g broth was taken and instantaneously quenched in pre-cooled pure methanol (−40°C), followed by a washing procedure with 80 % aqueous methanol (v/v) solution pre-cooled to −40°C. Metabolite extraction was performed with 75 % (v/v) ethanol (95°C, 3min), followed by rapid vacuum evaporation until dryness. A detailed protocol has been described previously (47). Metabolite concentrations were quantified by isotope dilution mass spectrometry (LC-IDMS/MS and GC-IDMS) using U-^13^C-labeled yeast cell extract as internal standard (52). Metabolites from glycolysis, TCA cycle and pentose-phosphate pathway as well as amino acids were quantified according to published protocols (53–55). Intracellular adenine nucleotide contents (ATP, ADP, AMP) were measured according to (55). The adenylate Energy Charge(AEC) was calculated as follows:

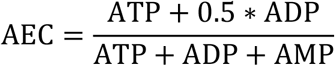

### Metabolic flux analysis

Intracellular flux distributions during steady-state chemostat and pseudo-steady-state retentostat cultivation were calculated using a slightly modified version of a previously published stoichiometric model (56), in which the biomass composition was adapted according to the measurements of the biomass elemental compositions. The input variables used for the flux analysis are summarized in supplementary Table S3.

## Acknowledgement

This research was financed by the Netherlands Be-Basic research program (Be-Basic project: FS10-04 Uncoupling of microbial growth and product formation).We thank Cor Ras, Patricia van Dam, Silvia Marine and Johan Knoll for analytical support.

